# Telomeric assemblies of *Paracoccidioides* genomes

**DOI:** 10.1101/2025.10.30.685552

**Authors:** Dmitry Grinevich, Oliver Kompathoum, Jingbaoyi Li, David A. Turissini, Patrick Kelly, McKenna Sutherland, Santiago Marin-Carvajal, Oscar M. Gomez, Gil Benard, Maria Aparecida Shikanai-Yasuda, Tiago Alexandre Cocio, Adriana Pardini Vicentini, Zoilo Pires de Camargo, Anderson Messias Rodrigues, Rosane Christine Hahn, Beatriz da Silva Motta, Priscila Marques de Macedo, Rodrigo de Almeida Paes, Primavera Alvarado, Rosely Maria Zancopé-Oliveira, Juan G. McEwen, Victoria E. Sepúlveda, Marcus M. Teixeira, Daniel R. Matute

## Abstract

*Paracoccidioides* is a genus of dimorphic fungal pathogens endemic to Latin America. We generated long-read de novo assemblies for 11 isolates representing four species of the *brasiliensis* complex (*P. brasiliensis, P. americana, P. restrepiensis, P. venezuelensis*) and *P. lutzii*. These include the first complete telomere-to-telomere assemblies for *P. brasiliensis* (Pb18) and *P. americana* (Pb03), each with five chromosomes. Comparative analyses revealed chromosomal fusion and fission events distinguishing *P. brasiliensis* and *P. americana*, and a 90 kb tandem duplication in *P. americana* containing siderophore biosynthesis genes (*sid1, sid3, sid4*), a cluster of putative virulence factors. Mitochondrial genomes showed conserved gene order but a phylogenetic topology inconsistent with the nuclear tree, suggesting mitochondrial introgression between *P. lutzii* and *P. venezuelensis*. RNA transposable elements were enriched near telomeres, correlated with genome size, and most abundant in *P. lutzii*. These assemblies provide key resources for understanding genome evolution and introgression in *Paracoccidioides*.

**SIGNIFICANCE:** Species of *Paracoccidioides* cause paracoccidioidomycosis, a systemic mycosis that remains a major public health problem in Latin America. Despite their clinical importance, genome evolution across the genus is poorly understood owing to the lack of complete reference assemblies. Here, we present the first telomere-to-telomere reference genomes for *P. brasiliensis* and *P. americana*, enabling a comprehensive comparison of chromosomal structure across the genus. Our analyses reveal that the nuclear genome is highly dynamic and shaped by large-scale rearrangements and structural variants, including the duplication of a siderophore biosynthesis-related gene cluster linked to virulence. In contrast, the mitochondrial genome is structurally conserved but shows introgression between species, revealing hidden evolutionary exchange. Together, these genomic resources redefine our understanding of *Paracoccidioides* evolution and provide a foundation for advances in molecular diagnostics, epidemiological surveillance, and studies of fungal pathogenicity.

## INTRODUCTION

Paracoccidioidomycosis (PCM) is a systemic fungal infection that primarily affects people in Latin America (Hahn et al. 2022; Rodrigues et al. 2023a). The disease is endemic to subtropical regions from Mexico to Argentina, particularly Brazil, Colombia, and Venezuela, and predominantly impacts agricultural workers in rural areas (Martinez 2015; Peçanha et al. 2022). PCM manifests in two main forms. The acute or subacute form involves the reticuloendothelial system and is more common in children and adolescents. The chronic form is characterized by chronic pulmonary symptoms, mucocutaneous lesions, and lymphadenopathy, often mimicking tuberculosis or malignancies. Other symptoms of chronic PCM include persistent cough, weight loss, and fever (Benard 2008; Restrepo et al. 2008; Ferreira 2009; Falcão et al. 2023). The chronic form primarily affects adult males aged 30–60, with a male-to-female ratio of 10:1. This disparity is attributed to higher environmental exposure in male workers, and the protective effects of estrogen against transition from mycelium to yeast (Restrepo et al. 1984; Shankar et al. 2011; Caixeta et al. 2018; Brito et al. 2021). The estimated annual incidence is 3–4 cases per 100,000 people per year in hyperendemic areas. Over 80% of cases occur in Brazil, where approximately 1–3% of all hospitalizations for systemic mycoses are attributed to PCM (Bellissimo-Rodrigues et al. 2010; Silva et al. 2021; Peçanha et al. 2022). Recent trends indicate shifting epidemiological dynamics of paracoccidioidomycosis, particularly in Rio de Janeiro, where outbreaks linked to environmental disturbances and an increase in PCM-related deaths suggest a concerning rise in disease burden (do Valle et al. 2017; Falcão et al. 2024). Treatment typically involves itraconazole or trimethoprim-sulfamethoxazole for mild cases, or amphotericin B for severe or disseminated forms (Visbal et al. 2005; do Carmo Silva et al. 2020).

The causal agent of PCM, *Paracoccidioides*, is a dimorphic fungus that transitions between yeast and filamentous hyphae depending on environmental conditions, mainly temperature. At environmental temperatures (25°C), *Paracoccidioides* grows in its mycelial saprophytic form, but at 37°C, it transitions to a pathogenic yeast form (San-Blas 1985; Borges-Walmsley et al. 2002; Theodoro et al. 2008). This temperature-mediated morphological switch is considered a key factor in the onset of virulence. The fungus thrives in riparian environments (Restrepo et al. 2001; Simões et al. 2004; Días and Ismael 2007) and has been associated with various terrestrial mammals (Richini-Pereira et al. 2008; Sbeghen et al. 2015; Losnak et al. 2018; Falcão et al. 2023), including armadillos (Bagagli et al. 1998; Corredor et al. 2005; Bagagli et al. 2021; Kluyber and Desbiez 2023). However, the ecological niche of *Paracoccidioides* remains largely unknown. Transmission to humans occurs primarily through inhalation of airborne conidia or mycelial fragments, which transition into the pathogenic yeast form after reaching the lungs. The latency period of PCM can range from months to decades (Benard 2008; Restrepo et al. 2008). Risk activities associated with increased exposure to *Paracoccidioides* include soil excavation, deforestation, handling of organic material, construction work, agricultural practices, and the building of cisterns or wells.

The genus *Paracoccidioides* was long considered to be monotypic. Early molecular genetic surveys revealed the existence of extensive genetic diversity across the continent, which eventually led to the recognition of species boundaries (Matute et al. 2006; Teixeira et al. 2009) and taxonomic reclassification (Teixeira et al. 2014; Turissini et al. 2017). Currently, the genus *Paracoccidioides* encompasses at least five distinct species, distributed across Latin America. The sister species *P. restrepiensis* and *P. venezuelensis* are primarily found in Colombia and Venezuela. *Paracoccidioides brasiliensis*, a sister species to this dyad, is found primarily across Brazil, Paraguay, and Argentina, and is the dominant species recovered from humans and animals. The most divergent member of the genus, *P. lutzii*, was confirmed as a distinct species through molecular analyses, which revealed differentiation from the rest of the *Paracoccidioides* species (Teixeira et al. 2009; Teixeira et al. 2014). All species pairs show low levels of gene flow (Muñoz et al. 2016; Mavengere et al. 2020) despite sympatry and the opportunity for admixture (Bagagli et al. 2021). Among the known species, *P. americana* and *P. lutzii* are the most distantly related, making them easier to distinguish through proteomic, molecular, and morphological assays (Turissini et al. 2017; de Oliveira et al. 2018; Cruz-Leite et al. 2023). Two more species, *P. lobogeorgii* and *P. cetii*, have been tentatively associated with the *Paracoccidioides* genus (Vilela et al. 2023; Vilela et al. 2023). However, their classification remains uncertain due to their unculturable nature.

Clinical and evolutionary studies have emphasized the need for well-assembled genomes across the entire *Paracoccidioides* genus (Hahn et al. 2022; Rodrigues et al. 2023b). To date, genomic research has focused primarily on two species, *P. brasiliensis* and *P. lutzii*. The genomes of *Paracoccidioides* species exhibit high levels of synteny, particularly between *P. brasiliensis* Pb18 and *P. americana* Pb03, with somewhat lower conservation in *P. lutzii* Pb01 (Desjardins et al. 2011). Despite this structural conservation, these genomes display notable gene family contractions, especially in carbohydrate-active enzymes, reflecting a reduced capacity for plant degradation compared to other filamentous fungi (Desjardins et al. 2011; Muñoz et al. 2014). In contrast, expansions have occurred in specific families—such as FunK1 fungal-specific kinases, proteases, and certain transposable elements—most prominently in *P. lutzii* (Desjardins et al. 2011). However, a comprehensive evaluation of synteny, chromosomal rearrangements, and other drivers of genome evolution in *Paracoccidioides* requires denser sampling across the genus phylogeny. In this study, we address this gap by examining genome evolution across the *Paracoccidioides* genus.

In this study, we expanded the use of long-read sequencing to generate *de novo* assemblies of highly contiguous genomes for eleven isolates representing five *Paracoccidioides* species. Two of these genomes (*P. brasiliensis* and *P. americana*) were assembled at the complete chromosomal level, while the remaining nine assemblies are also highly contiguous. We found that transposable element content varies both within and between species. Although overall gene content and genome organization are relatively consistent, we identified large-scale chromosomal rearrangements that have occurred during the evolution of *Paracoccidioides*. Our characterization of genome architecture in these human fungal pathogens reveals dynamic evolution in both content and structure.

## MATERIALS AND METHODS

### *Paracoccidioides* isolates and DNA extraction

The *Paracoccidioides* strains included in this study are listed in Table S1. We cultivated *Paracoccidioides* yeast cells on BHI and Fava-Netto agar slants at 37°C, subculturing them every 15 days. We harvested approximately 1g of yeast biomass and lysed the yeast cells with protocols using maceration of frozen cells, or glass beads (Van Burik et al. 1998). We performed High Molecular Weight (HMW) DNA extraction using the Quick-DNA™ HMW MagBead Kit (Zymo Research), following the manufacturer’s instructions. To assess DNA integrity and concentration, we ran the samples on an 1% agarose gel with a DNA mass ladder, visualizing it with ethidium bromide and UV light, and performed spectrophotometric analysis using a NanoDrop™ 2000 (Thermo Scientific).

### Genome sequencing

Each isolate was sequenced using Oxford Nanopore technology (Abingdon, UK). Libraries were barcoded using the Native Barcoding Kit 96 (SQK-NBD112.96), following the manufacturer’s protocol and sequenced on r10.4.1 chemistry flow cells. Basecalling was performed with Dorado v4.2.0 at 400 bp using the super accurate model. Sequencing was carried out on a MinION Mk1B device using MinKNOW software v23.07.12 (Oxford Nanopore). Each run lasted 72 hours and generated POD5 files. Table S1 includes total read counts, estimated base qualities, and SRA accession numbers.

### Assembly Pipeline

We used FASTQC version 0.12.1 (available at https://www.bioinformatics.babraham.ac.uk/projects/fastqc/) to estimate quality control metrics (e.g. read size, N50 values, read quality). We performed *de novo* genome assembly using raw Nanopore reads with Canu version 2.2 (Koren et al. 2017). We used the default Canu command with the parameters (genomeSize=40m, gridOptions=”—time=72:00:00”, -nanopore). The assembly results are shown in Table S2. Next, we polished the assembly using Pilon version 1.24 (Walker et al. 2014). Polishing uses existing read sequencing data aligned to the draft reference genome and corrects the draft genome based on evidence supported in reads to fix base errors, fill gaps, and correct mis-assemblies (Walker et al. 2014). We used bwa-mem2 version 2.1.1 (Vasimuddin et al. 2019) to generate the aligned SAM format files required for polishing. We generated genome indices using BWA index, employing 24 threads (-t 24) and the long-read setting optimized for Oxford Nanopore data (-x ont2d). Next, we used Samtools version 1.1 (Li et al. 2009) to generate an index file. Finally, we performed three rounds of Pilon polishing for each *Paracoccidioides* assembly. Telomeres were identified by manual inspection of each contig, searching for 5’-TAACCC-3’ and 3’-GGGTTA-5’ on each corresponding contig end. We required at least three repeats of the telomeric motif to consider the sequence telomeric. Assemblies for which we detected all telomeres were denominated as chromosomal. Assemblies missing more than one telomere were called highly contiguous. Next, we assessed the completeness of the assembled genomes by analyzing the percentages of orthologs present in each assembly. To this end, we use single-copy orthologous genes. We extracted coding sequences from 3,478 genes identified in the Eurotiomycetes dataset of the BUSCO (Benchmarking Universal Single-Copy Orthologs) ortholog database, version odb10 (Waterhouse et al. 2018; Kriventseva et al. 2019; Manni et al. 2021). The Eurotiomycetes dataset provides a curated set of orthologous genes expected to be present as single copies in most species within this fungal class. We used BUSCO tool version 5.5 to perform analyses (Simão et al. 2015; Waterhouse et al. 2018).

### Gene Prediction and Functional Annotation

We performed gene prediction in the assembled genomes described above using a combination of evidence-based and *ab initio* approaches implemented in the *funannotate* pipeline (version 1.8.17; (Palmer 2017) available at GitHub: https://github.com/nextgenusfs/funannotate). We removed duplicate contigs using minimap2 v2.28-r1209 (Li 2018) through the *funannotate clean call*, sorting by smallest contigs and removing duplicates. Contigs were then reordered from longest to shortest, and FASTA headers were renamed for clarity using *funannotate* sort. Reads were softmasked using Tantan v49 (Frith 2011) as implemented in *funannotate mask*. Assemblies were then piped into *funannotate predict*, which included multiple *ab initio* gene prediction tools. GeneMark-ES v4 (Lomsadze et al. 2005) was then used in self-training mode to predict protein-coding genes. BUSCO v2 was used to collect conserved gene models for prediction (BUSCO v2 was used in funannotate due to pipeline compatibility). To guide the identification of conserved gene structures, we used Augustus v3.5.0 (trained with *Histoplasma* sp. gene models; (Stanke and Morgenstern 2005)). Additional gene predictions were conducted using SNAP and glimmerHMM v3.0.4 (Korf 2004; Majoros et al. 2004). To integrate predictions and generate a consensus gene model, we utilized EVidenceModeler (Haas et al. 2008) with the following weights: Augustus = 1, HiQ= 2, GeneMark= 1, GlimmerHMM= 1, snap= 1. All other protein models were weighted at 1. Protein FASTAs were generated from EVidence Modeler, and inaccurate gene models were filtered using diamond v2.1.9. tRNAs were predicted using tRNAscan-SE v2.0.12 (Chan et al. 2021). Finally, gene annotations were converted to Genbank format and saved for each isolate.

Next, we analyzed the contraction and expansion of orthologous gene families across all analyzed isolates within our dataset. We built related ortholog cluster groups using the *funannotate* compare module (Palmer 2017). We focused on three gene classes: the transcription factors present in the InterPRO database (Paysan-Lafosse et al. 2023; Blum et al. 2025), the Carbohydrate-Active enZYmes database (CAZy, (Cantarel et al. 2009)), and the MEROPS peptidase database (Rawlings et al. 2010). Consensus gene predictions were consolidated and exported in GenBank format (.gbk) for each isolate using the annotate function of *funannotate*.

To determine whether observed copy numbers deviated from neutral expectations along the phylogeny, we applied CAFE5 (Mendes et al. 2020), which uses maximum-likelihood to estimate gene family evolutionary rates. CAFE5 requires an ultrametric tree. We rooted with *Histoplasma* Africa H88 using the R function *reroot* (package *ape*, (Paradis and Schliep 2019)) the nuclear genome derived phylogeny (see Phylogenetic Tree Inference below), then the make_ultrametric.py script from the Orthofinder suite v2.5.5 (Emms and Kelly 2015; Emms and Kelly 2019). We scaled the branch lengths of the tree to reflect the tip-to-tip estimate age between *P. lutzii* and the *brasiliensis* species complex (∼30 million years diverged, (Teixeira et al. 2009)). We ran CAFE5 with default parameters.

### Synteny

To show the patterns of genome evolution in *Paracoccidioides* we used synteny plots. We included the 11 assemblies we generated in this report, one long-read (ONT) based assembly of Pb369 (Lorenzini Campos et al. 2024), and three assemblies based on fasmids and paired-end shotgun Sanger sequencing (*Paracoccidioides lutzii* Pb01 (ABKH00000000), *Paracoccidioides americana* Pb03 (ABHV00000000), and *P. brasiliensis* Pb18 (ABKI00000000; (Desjardins et al. 2011)). We conducted comparative genomic analyses using GENESPACE v1.1.4; (Lovell et al. 2022), a pipeline designed to identify syntenic gene orthologs and assess gene family dynamics across multiple genomes. BED file positions were extracted from associated general feature format (GFF3) files from genome annotations. Annotated genome assemblies in GFF3 format and corresponding protein FASTA files were used as input. GENESPACE was run with default parameters using MCScanX (Wang et al. 2012) for synteny block detection, and OrthoFinder 2.5.5 (Emms and Kelly 2019) for gene family clustering. We specified the following parameters for OrthoFinder: -t 6, -a 1, -X. To anchor syntenic comparisons, a reference genome was specified, and orthologous gene relationships were inferred across all genomes. The resulting gene family presence–absence matrix and synteny plots were used to visualize gene gain/loss patterns and structural rearrangements. Downstream analyses included filtering for gene families with lineage-specific expansions and mapping their genomic locations relative to conserved syntenic blocks. We used the following command line: gpar <- run_genespace(gsParam = gpar) to generate initial Genespace files and for plotting we used ripDat <- plot_riparian(gsParam = gpar, refGenome = “Pb18”, forceRecalcBlocks=FALSE, invertTheseChrs = invchr). Chromosomes were inverted if required for visual clarity, and the Pb18 genome was used as a reference to set synteny colors and blocks. All our scripts for this portion of the research are available at GitHub (https://github.com/Dogrinev/).

### Structural Variant Analysis

Next, we investigated structural variation across all *Paracoccidioides* isolates. Structural variants were identified with SVIM v2.0.0 (Heller and Vingron 2019) using the standard command line. We performed variant calling with reference-read alignments against both the *P. brasiliensis* and *P. americana* complete genomes, which served as alternative reference genomes. Variant call format (VCF) files were generated for each mapping–reference combination and filtered with a minimum support cutoff of 10 to exclude low-confidence structural variants. Output GFF3 files were then used to identify the start and stop positions of each detected variant across the *Paracoccidioides* genomes.

To assess genomic distance and evaluate potential reference bias, we applied the MinHash dimensionality-reduction approach (Mash available at https://github.com/marbl/mash (Ondov et al. 2016). Pairwise genome comparisons were performed using the *mash dist* command with standard parameters, and for each genome pair we recorded both the Mash distance and the number of matching hashes.

Given the potential impact of reference bias (Ballouz et al. 2019; Günther and Nettelblad 2019), we assessed how reference choice influences structural variant detection. Specifically, we mapped reads from a *P. brasiliensis* isolate (113) to both the complete *P. brasiliensis* reference genome (Pb18) and the complete *P. americana* reference genome (Pb03), both generated in this study. We also performed the reciprocal analysis by aligning reads from a *P. americana* isolate (28434) to both reference genomes (Pb18 and Pb03).

SVIM also identifies structural variants and assigns them a quality score between 0 and 100 to represent the confidence of the detected structural variant. Quality scores (S) are represented through the equation:

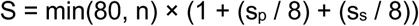

Where (n) is the number of “signatures” or supporting reads for a detected variant, (s_p_) is the standard deviation of the *positions* of the signatures in a cluster, normalized to the average span length of reads in the cluster, and (s_s_) is the standard deviation of the span of the signatures in a cluster, normalized to the average span length of reads in the cluster (Heller and Vingron 2019). For this study, we considered putative structural variants to be high confidence if their score was ≥ 90.

### mtDNA assembly

To identify putative mtDNA contigs, we scanned each genome assembly FASTA file for contigs with elevated coverage (>10× higher than nuclear DNA contigs). As additional support, we required that each putative mtDNA contig was flagged as *sugCirc=TRUE* in the Canu *contigs.layout.tigInfo f*ile, indicating circularity. We then used BLAST to confirm the presence of three mitochondrial genes (*cox1*, *atp6*, and *nad5*) within each candidate mtDNA contig. For *Pb339*, the mitochondrial sequence was reconstructed from two contigs in the original Canu assembly. Both contigs were aligned to the Pb18 mitochondrial sequence using BLASTN. Overlapping regions between contig 406 and contig 410 were manually recorded, trimmed with *samtools faidx* (Li et al. 2009), and the unique sequences from each contig were combined to generate the complete mtDNA sequence for *Pb339*.

After confirmation of mtDNA contigs for each isolate, we used *MFannot* (Lang et al. 2023) to annotate mtDNA genes. We used the yeast mitochondrial genetic code setting for gene prediction. We manually converted the TBL files provided by *MFannot* into SEQ table files. We checked all annotation tables and mitochondrial sequences for the presence of *rns* and *rnl* in the cases in which they were not detected by *MFannot*. In those instances, we used *BLASTN* and manually inserted the gene with ranges provided by the blast hits if they were missing. We individually uploaded all SEQ files into *circularMT* and manually adjusted them such that each graph was anchored to the first putative copy of *nad2*. We visualized the annotated mitochondrial sequences using *CircularMT* (Goodman and Carr 2024).

### Genomic and Mitochondrial sequence Dotplots

We did several pairwise comparisons between contigs. To do so, we constructed dotplots for both the genomic and mitochondrial sequences of *Paracoccidioides* using the dotPlotly R script (Poorten 2017) to compare base pair similarities between isolates. First, we used minimap2 v (Li 2018) to generate assembly-to-assembly mappings in PAF format. We generated the PAF files using the asm5 preset, which is preferred for aligning long assemblies to a reference with low sequence divergence. We input the PAF files into the dotPlotly R script with the minimum alignment length and minimum query length set to 1 so that the mitochondrial sequences could be recognized. For visualization, we concatenated the image files into a PDF using imagemagick v7.1.1-47 (The Definitive Guide to ImageMagick 2006).

### Phylogenetic Tree Inference

We generated phylogenetic trees to understand the different evolutionary histories of the nuclear and the mitochondrial genome. We describe the three steps for these analyses as follows.

#### Nuclear genome-based tree

We inferred a phylogenetic tree on all assembled *Paracoccidioides* genomes in this study using the Universal Fungal Core Genes (UFCG) analysis suite (Kim et al. 2023). Assemblies of our genomes were input in FASTA format to UFCG profile, which utilized the BUSCO fungi_odb10 to extract a set of 758 core and marker genes from across fungal taxa. Using the extracted gene sequences, we generated multiple sequence alignments for each gene using *MAFFT* v7.310 (Katoh and Standley 2013; Katoh and Standley 2014) which were then concatenated into a single file. We removed characters in the supermatrix alignment that were over 50% missing. We enabled the Modelfinder (Kalyaanamoorthy et al. 2017) module in IQTREE 2, which found that the GTR+F+R4 substitution model was the best model fit. Next, we used the UFCG tree function with optional parameters (-t 10, -p iqtree, -l strain). We used the *IQ-TREE* v2.3.6 to infer a maximum-likelihood tree. We rooted the tree using *H. capsulatum* isolate H88 as an outgroup. We estimated branch support using the built-in IQ-TREE 2 module and 1,000 replicates of ultrafast bootstrapping.

#### mtDNA genome-based tree

We used MAFFT v7.526 (Katoh et al. 2019) to generate a multiple sequence alignment (MSA) of the mtDNA sequences. Similar to the nuclear tree, we used *H. capsulatum var. duboisii* (H88) as an outgroup. Once the MSA was generated, we ran IQTREE 2 (Minh et al. 2020), including the Modelfinder (Kalyaanamoorthy et al. 2017) module, to generate a maximum likelihood tree from the aligned mtDNA sequences. We used the same scheme of bootstrapping as described for the nuclear phylogeny to estimate branch support.

#### Tree comparison

We compared the topologies of the nucDNA and mtDNA trees using a Robinson-Foulds (RF) distance (Bryant and Steel 2009) using the R function *treedist* (library *phangorn*, (Schliep 2011)). This index ranges between 0 (no concordance between trees) and 1 (complete concordance between trees). Additionally, we forced the two trees to become ultrametric using the function *force.ultrametric* (library *phytools,* (Revell 2012; Revell and Revell 2014)) and calculated the root-to-tip branch of the two trees.

### Transposable Element Analysis

Transposable elements (TEs) are key drivers of genome evolution dynamics (Kidwell 2002; Grandaubert et al. 2014; Krasileva 2019; Muszewska et al. 2019). We studied the TE content in *Paracoccidioides* using the long-read assemblies described above. First, we identified TEs by running EDTA v2.2.0 (Ou et al. 2019) to generate a *de novo* TE library for each genome. We then used RepeatMasker v4.1.5 (Chen 2004) to search and annotate TEs in each genome based on these TE libraries. To process the RepeatMasker output, we calculated the total length of each TE class and normalized these values by genome size, yielding the proportion of the genome occupied by each TE category (code available in GitHub: https://github.com/Dogrinev/).

We studied two hypotheses with this data. First, we evaluated whether transposable element (TE) content correlates with genome size by fitting regressions in R, using both ordinary least squares (OLS) and phylogenetic generalized least squares (PGLS). We used the R packages *ape* (Paradis and Schliep 2019), *nlme* (Pinheiro et al. 2017), *geiger* (Harmon et al. 2008; Pennell et al. 2014; Harmon et al. 2015), and *phytools* (Revell 2012) for PGLS; the R package *emmeans* for model inference; and the R package *effects* for extracting model parameters. We performed these analyses both with and without the outgroup, *Histoplasma*, which showed the largest genome size among our focal taxa. We also visualized TE content across the phylogeny of *Paracoccidioides*. We used the R package *ggtree* v3.10.0 (Yu et al. 2017) to generate a phylogenetic tree with stacked bar plots representing the genomic proportions of each TE class.

Second, we studied whether the extent of genetic differentiation in TEs paralleled that of the rest of the genome using Principal Component Analysis (PCA). First, we constructed a variant call file (VCF) of the whole genome and filtered sites using the *bcftools mpileup* and *bcftools* call commands (Li 2011; Danecek et al. 2021), annotating for allelic depth, genotype depth, and strand bias and formatted for genotype quality and genotype probability. We used PLINK (Purcell et al. 2007; Chang et al. 2015) to filter the VCF to only include biallelic sites with minor allele frequency greater than 0.05, and to exclude sites with missing call rates. We also used PLINK to estimate the covariance matrix among sites and visualize the principal components. We plotted the combinations of PC1/PC2 and PC3/PC4. We evaluated the contribution of each PC using a Scree plot. Next, we performed a more limited PCA analysis on TE-specific regions by subsetting our *Paracoccidioides* genome VCF. Since we prepared the VCF initially by aligning all isolates to our complete Pb18 genome, we used the identified TE locations from EDTA to subset the VCF to include only TE sites. For this second PCA, we used an identical approach as the one described immediately above for the genome wide variation.

## RESULTS

### The *Paracoccidioides* nuclear genomes are largely but not completely syntenic

We sequenced and assembled 11 isolates of *Paracoccidioides* using long-read Nanopore sequencing data from the five culturable species of *Paracoccidioides*: *P. brasiliensis*, *P. americana*, *P. restrepiensis*, *P. venezuelensis*, and *P. lutzii*. Of these 11, sequencing of two isolates, Pb18 (*P. brasiliensis*) and Pb03 (*P. americana*) yielded chromosomal-level assemblies. The coverage of these two genomes was 62.7x and 100x read coverage, respectively. In both cases, the genome assembly encompassed five complete chromosomes and a single closed mitochondrial genome. For both genomes, we identified a total of six contigs each (Figure 1A). Five of these contigs corresponded to the complete chromosomes, while the sixth represented the circular mitochondrial DNA segment (see below). All chromosomal contigs were confirmed to have telomeric repeat caps (5’-TAACCC and 3’-GGGTTA).

**FIGURE 1.**
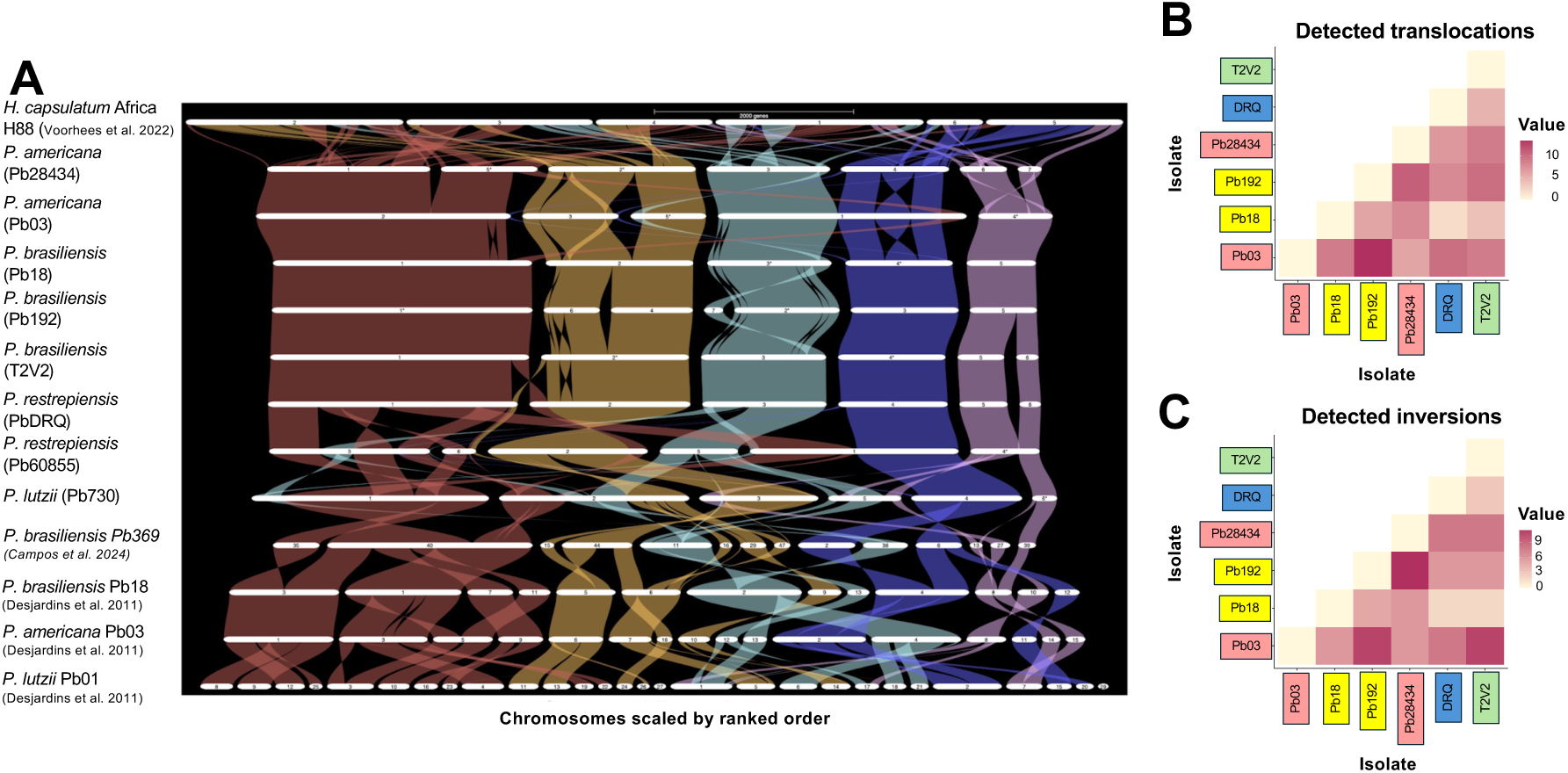
*Paracoccidioides* Genome Synteny. Syntenic analysis using GENESPACE of telomeric and highly contiguous *Paracoccidioides* genomes included in this study and previous studies. Complete telomere to telomere assemblies are indicated in bold and red label text. Assemblies generated in previous studies are included with the original data source. The complete synteny includes *P. brasiliensis, P. americana, P. venezuelensis, P. restrepiensis, and P. lutzii.* A significant chromosomal exchange happens between complete genomes of *P. brasiliensis* and *P. americana.* While the primary chromosome (indicated in red) is conserved across species, the smaller chromosomes (blue and teal) combine into a single secondary large chromosome in *P. americana*. *P. restrepiensis* shows the most differences in chromosomal changes, including exchange between the primary chromosome (red). Our syntenic analysis confirms that partially complete assemblies have gaps in chromosomes which are parts of an un-assembled larger chromosome. **B.** Pairwise comparisons between genomes showing the number of translocations. **C.** Pairwise comparisons between genomes showing the number of translocations. For B and C, only the top six assemblies are shown.

In addition to these two complete assemblies, we assembled six highly contiguous genomes (defined as assemblies with at least 3+ complete telomere-capped chromosomes). This group encompassed two isolates of *P. restrepiensis* (PbDRQ, and Pb60855) and single isolates of *P. brasiliensis* (Pb192)*, P. americana* (28434)*, P. venezuelensis* (T2V2), and *P. lutzii* (7730). Finally, we generated partially-complete assemblies (defined as assemblies with 0-2 complete chromosomes) for one *P. restrepiensis* isolate (Pb339), one *P. brasiliensis* isolate (Pb113), and one *P. lutzii* isolate (Pb01). Table S2 shows the summary statistics for each of these assemblies. All the assemblies were deposited in SRA (Accession numbers: TBD).

We used these genome assemblies to study the extent of synteny within the *Paracoccidioides* genus. Figure 1 shows 8 of the 11 genomes we generated, along with all previous *Paracoccidioides* assemblies (Desjardins et al. 2011; Lorenzini Campos et al. 2024), and an isolate of *Histoplasma capsulatum Africa* H88 (Mapengo et al. 2025), a related species of dimorphic fungi. We excluded three genomes from the figure (Pb339, Pb01, and Pb113) due to lower assembly quality in these three isolates. Synteny assessments revealed that the two largest chromosomes are largely conserved in size and structure across species in the genus with approximately 10 and 6 Mb (Fig 1, colored red and orange). This approach also revealed that the near-complete assembly of *P. americana,* which contains six contigs, also has five chromosomes as the two smallest two contigs are two pieces likely to be part of the same chromosome (Figure 1, purple). Other chromosomal rearrangements seem to have a biological rather than technical explanation. For example, compared to all the other *Paracoccidioides* genomes, the complete *P. americana* genome, Pb03, chromosome 2 is split into two telomerically capped smaller chromosomes.

A notable example of the fairly conserved structure of the genome of *Paracoccidioides* within the *brasiliensis* species complex was the line Pb60855 (*P. restrepiensis*). This clinical isolate was collected from a patient and has been kept in culture for over 45 years (Restrepo-Moreno and Schneidau 1967). A maximum likelihood phylogenetic tree confirmed that the line is, indeed, part of *P. restrepiensis* (starred in Figure 3A) thus ruling out contamination. We generated genomic alignment dotplots of Pb60855 to study its structure compared to related *Paracoccidioides* isolates. We generated five pairwise comparisons, with dotplots showing genomic alignment between Pb60855 and four other isolates from four different species: *H. Africa* H88, Pb18, Pb03, and Pb7730 (Figure 2). Dissimilarity in gene order is higher as genetic distance increases. As expected, a second isolate of *P. restrepiensis*–DRQ– is the isolate with the highest level of gene order similarity to *P. restrepiensis* Pb60855 (Figure 2). As expected, Pb60855 and H88 differ significantly and show little genomic contiguity due to the genetic divergence between *Histoplasma* and *Paracoccidioides*.

**FIGURE 2.**
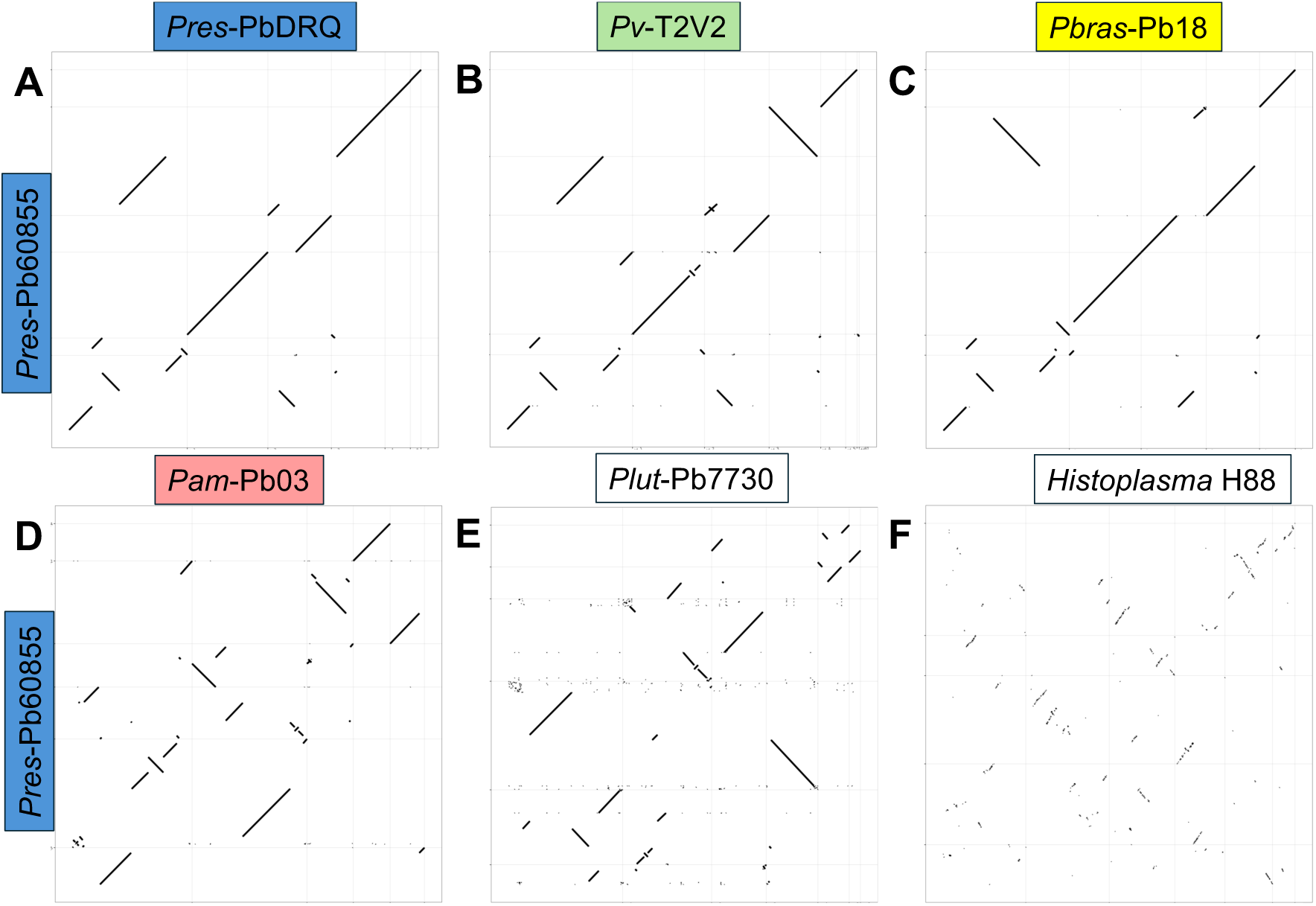
Dotplots comparing *P. restrepiensis* Pb60855 with other isolates of *Histoplasma* and *Paracoccidioides*. Each panel shows the sequence pairwise comparison (dotplot) between Pb60855 and a second isolate. **A.** *Histoplasma Africa* H88. **B.** *P. lutzii* Pb7730. **C.** *P. americana* Pb03. **D.** *P. brasiliensis* Pb18. **E.** *P. restrepiensis* DRQ.

**FIGURE 3.**
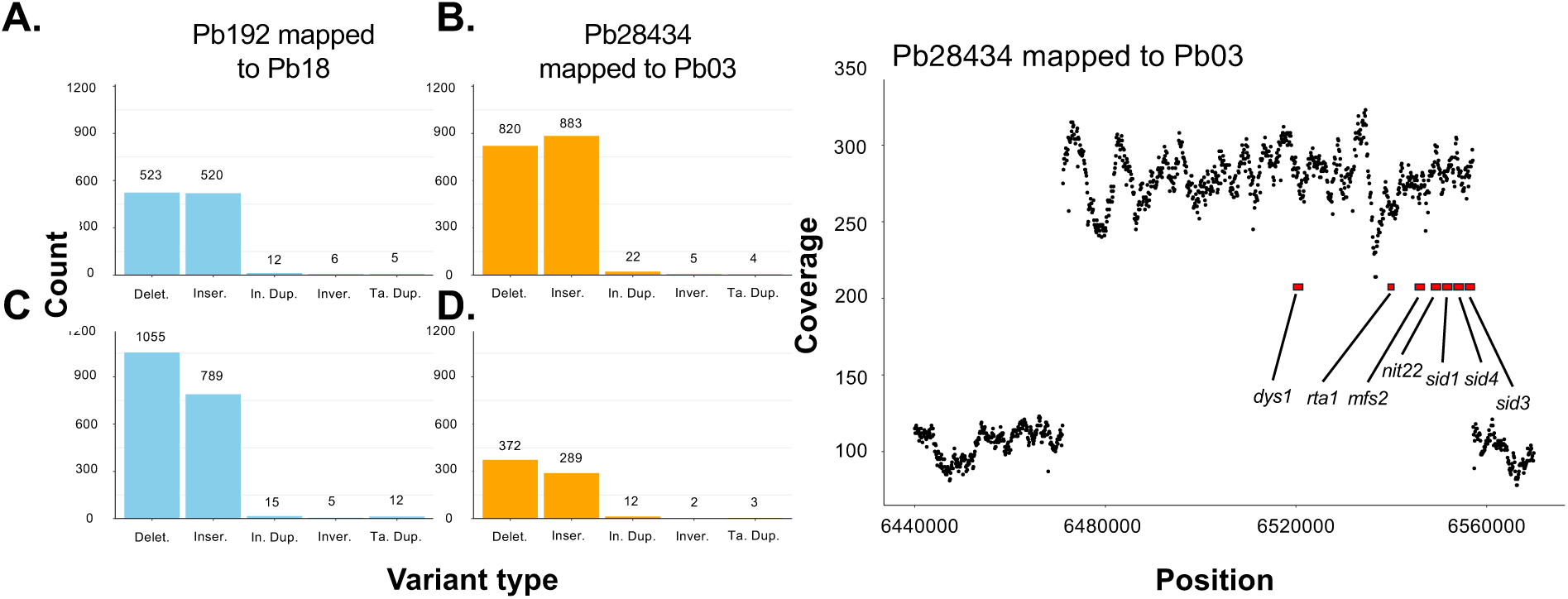
Reference bias approaches indicate the existence of structural variants within species of *Paracoccidioides*. **A-D.** Structural variants (SVs) were mapped to two different complete reference genomes available in our study (*P. brasiliensis* Pb18 and *P. americana* Pb03). In this example, we mapped *P. brasiliensis* isolate 113 and *P. americana* isolate 28434 to each reference genome to evaluate bias effects. We observed large changes in SV detection depending on the reference genome selected, with a bias towards certain types of SVs. We found that insertions, deletions, and duplications are heavily affected by reference choice. Other SV types, including inversions, tandem duplications, and breakends appear unaffected by reference choice. **E**. The only tandem duplication with strong support in SVIM shows an abnormally high degree of coverage with respect to the rest of the genome.

Next, we used the genome alignment to identify the relative abundance of chromosomal rearrangements. First, we focused on counting the number of rearrangements within the *brasiliensis* species complex. Among the highly contiguous and near-complete or complete assemblies (N=6 isolates), we identified a total of 236 translocation events and 201 inversion events (Table S3). Among these events, we detected 10 translocations and 11 inversions that were larger than 1 million base pairs. The largest inversion event we detected was a ∼1.12 million base pair inversion which occurs in chromosome 1 in one isolate of *P. americana* (isolate Pb03; Figure 1, blue). Of note, this inversion is not present in a second isolate of *P. americana* (28434) or any of the other *Paracoccidioides* isolates.

Large inversion and translocation events were most common in *P. americana* (Pb03 and 28434 strains) and *P. restrepiensis* (Pb-DRQ), showing 11, 7, and 9 total events relative to other strains (Table S3); *P. brasiliensis* (Pb18 and 192 strains) and *P. venezuelensis* (T2V2),had lower number of translocations in with 6, 6, and 5 events relative to other strains, respectively.

Species within the *brasiliensis* complex share a more similar genome structure with one another than with the more distantly related *P. lutzii*. The latter exhibits pronounced structural differences compared to other *Paracoccidioides* species. In particular, *P. lutzii* isolate 7730 shows extensive chromosomal rearrangements relative to other members of the genus. Although some chromosomes in *P. lutzii* (1, 3, and 4) appear relatively—but not entirely— conserved, the overall genome architecture of *P. lutzii* is distinct from that of the other *Paracoccidioides* species.

Next, we used a reference-based method to detect chromosomal variant breakpoints. Reference selection has large effects on the observed number of insertions, deletions, and duplications (Figure 3, Table S4). The structural variant count increased with the genetic distance between the reference genome and the mapped isolates which could be explained by genetic divergence, or by reference bias (Figure S1). To differentiate between these two possibilities, we mapped *P. brasiliensis* and *P. americana* reads to reference genomes of their own species and to reference genomes of the reciprocal species. The choice of reference bias had a strong effect on the detected number of deletions and insertions in these two species (Figure 3A-D). Similarly, the inferred number of duplications are affected by reference choice but to a lesser extent (Figure 3A-D). Given this reference genome biases, we focused on detecting duplication events within two of the species of *Paracoccidioides*, *P. brasiliensis* and *P. americana*.

We analyzed the presence of tandem and interspersed duplications within two *Paracoccidioides* species. We mapped seven duplication events in *P. brasiliensis* Pb113, 12 duplication events in *P. brasiliensis* strain 192, and 12 duplication events in *P. americana* strain 28434. We found two tandem duplications in *P. brasiliensis* Pb113, five tandem duplications in *P. brasiliensis* strain 192, and three tandem duplications in *P. americana strain* 28434. All putative duplication events are listed in Table S4. The majority of these events (40 out of 41) have moderate quality scores (S ranged between 10 and 25). One duplication in *P. brasiliensis* strain 113 showed strong support (S = 99), with respect to the other *P. americana* line, 28434 (Figure 3). The length of this tandem duplication was about 90kb and encompassed six genes: *rta1*, *mfs2*, *nit22*, *sid1*, *sid4*, and *sid3*. As expected, this region shows an abnormally high sequencing coverage (Figure 3C).

### The mtDNA gene order is highly conserved in *Paracoccidioides*

We also assembled the mitochondrial genome of *Paracoccidioides* for the 11 sequenced *Paracoccidioides* isolates. The initially inferred *Paracoccidioides* mtDNA genomes ranged from 130 Kb to 186 Kb. These assemblies suggested that three isolates *P. americana* (Pb03 and Pb28434) and *P. venezuelensis* (T2Z2) had a duplication of the mtDNA genome (Figure S2A). Sequence comparisons revealed that the two putative copies were identical, which in turn suggests this is not a true duplication. After sequence curation (Figure 4 *cf.* Figure S2), the mitochondrial genomes showed a consistent size of 115 Kb (± 2.6 Kb), which is in agreement with previous reports for *Paracoccidioides* mtDNA (Cardoso et al. 2007; Misas et al. 2020). All isolates had all ATP synthase genes (*atp6*, *atp8*, *atp9*), cytochrome associated genes (*cob*, *cox1*, *cox2*, *cox3*), NADH dehydrogenase associated genes (*nad1*, *nad2*, *nad3*, *nad4*, *nad4l*, *nad5*, *nad6*), and a full set of tRNAs required for the mitochondria (Figure 4). (The annotation pipeline did not consistently identify the presence and exact location of the small and large ribosomal subunits–*rnl* and *rns*–nonetheless, manual inspection identified the genes.) A comparison of the gene order in the eleven genomes revealed that the mtDNA gene order is identical in all isolates of *Paracoccidioides* (Figure 4, Figure S3), thus revealing conservation across the whole genus.

**FIGURE 4.**
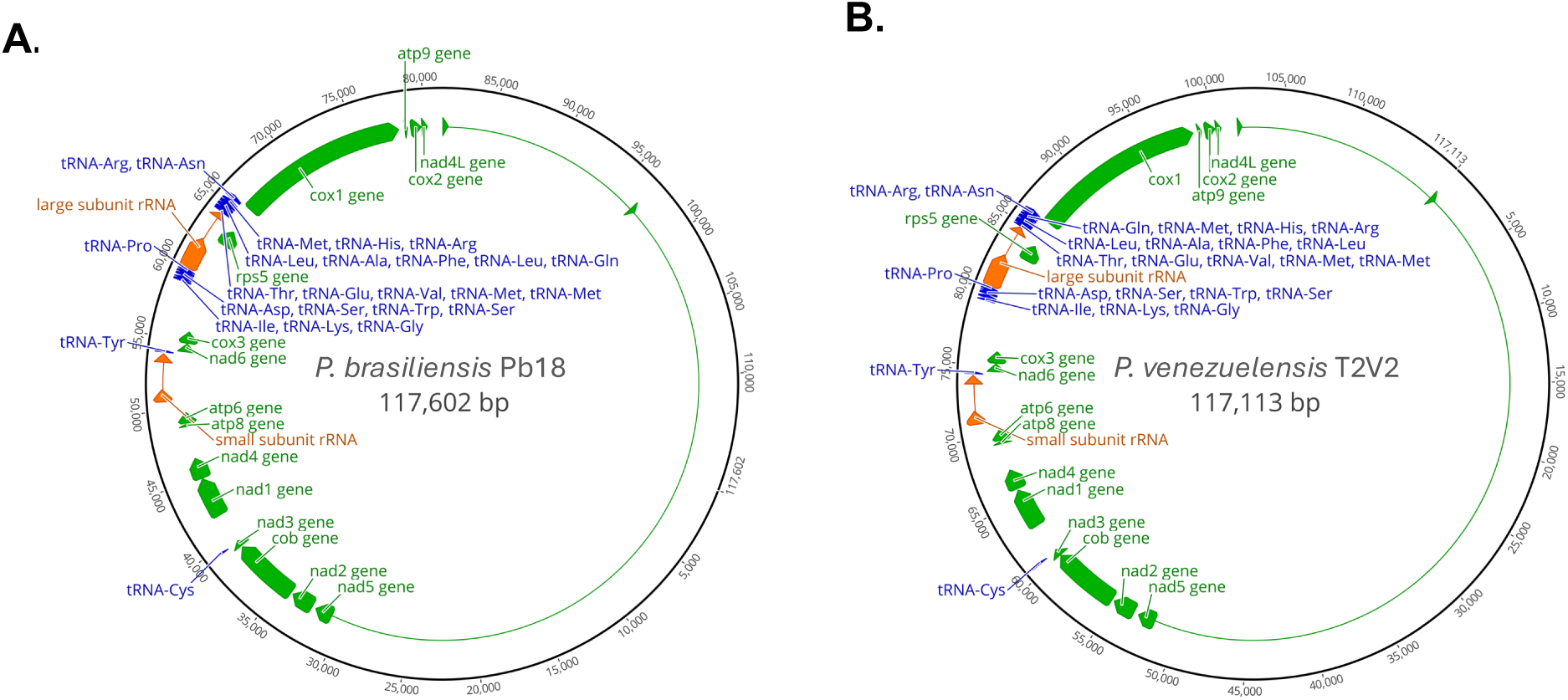
High degree of conservation in the mtDNA genome in the genus *Paracoccidioides*. **A.** Mitochondrial genome of *P. brasiliensis* Pb18. **B.** Mitochondrial genome of *P. venezuelensis.*T2V2. The comparison of the two genomes reveals that gene order is highly conserved across the whole genus.

### The *Paracoccidioides* nuclear and mitochondrial genomes show different phylogenetic histories

Next, we generated phylogenetic trees for the 11 samples of *Paracoccidioides* using the nuclear and mitochondrial genomes. The nuclear ML tree is in agreement with trees generated from unassembled genomes (Mavengere et al. 2020; Bagagli et al. 2021) and reflects the proposed species relationships between the species of the *brasiliensis* complex and of the group with *P. lutzii*. All the previously inferred reciprocal relationships with multilocus sequence typing (Matute et al. 2006; Teixeira et al. 2009b; Turissini et al. 2017) or unassembled genomes, are recapitulated with phylogenetic analyses with long-read assemblies (Figure 5A).

**FIGURE 5.**
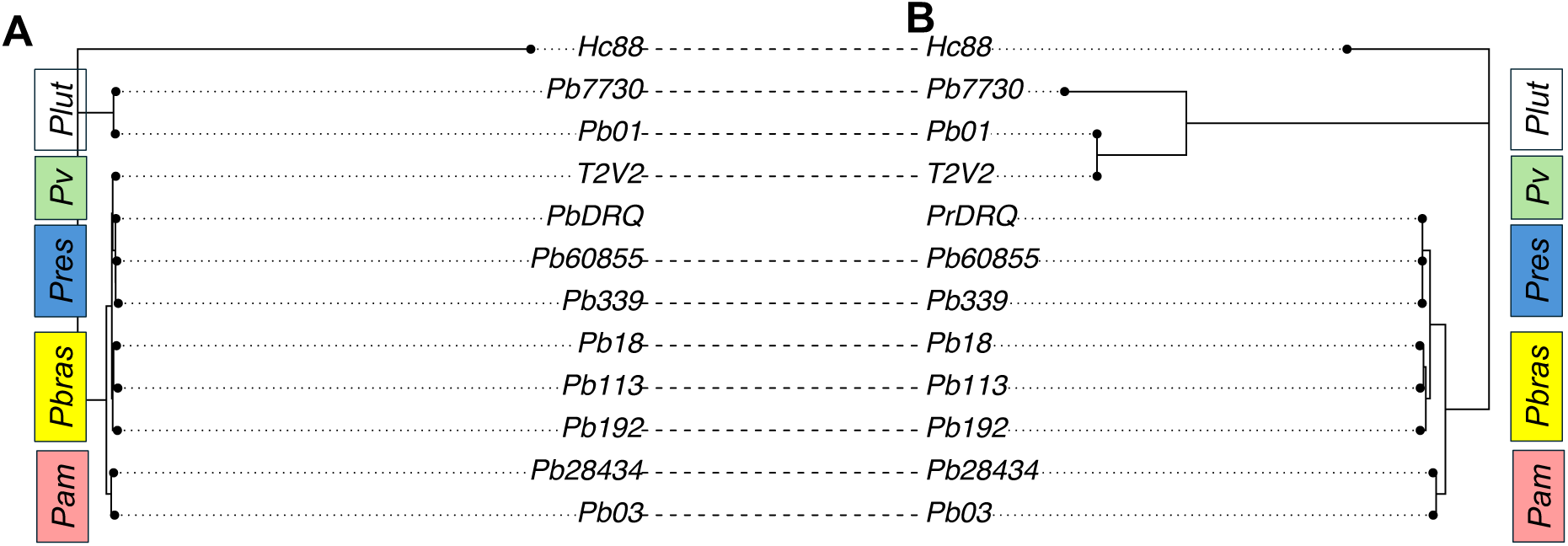
Phylogenetic trees. **A.** nucDNA**. B.** mtDNA. Connecting lines between the trees are meant to show instances of tree discordance. The main difference between the two trees is the position of *P. venezuelensis*.

In parallel, we generated a rooted tree for the mtDNA genome. While three of the species of the *brasiliensis* group show reciprocal relationships similar to that obtained in the nucDNA tree, *P. venezuelensis* appear nested within *P. lutzii* (Figure 5B). Consequently, the phylogenetic discordance between the two trees is high (Normalized RF distance = 0.444). This inconsistency between the nuclear gene genealogies and mtDNA loci was already reported with smaller genetic sampling (five mtDNA loci, (Salgado-Salazar et al. 2010; Turissini et al. 2017)). Our results indicate that the discordance extends to genome-wide markers. The root-to-tip length of the mtDNA tree was about 4-times larger than that of the nuclear tree. This tree length ratio is considerably lower than expected given the differences in mutation rate between mtDNA and nuclear DNA in yeast (∼11X, (Liu and Zhang 2019)). We thus hypothesize that mtDNA in *Paracoccidioides* follows a similar pattern to other organisms in which it can move between species much more readily than the nuclear DNA, and that the short branches are caused by interspecific introgression, a phenomenon that is almost absent in the nuclear genome (Mavengere et al. 2020).

### Transposable Elements in the *Paracoccidioides* genome

Transposable elements are considered an important driver of genome evolution in some, but not all taxa (Kidwell 2002; Lee and Kim 2014; Marino et al. 2025). Genetic surveys have revealed that the *Paracoccidioides* genome harbors TEs (Marini et al. 2010; Alves et al. 2014; Soares et al. 2015). We quantified TE content variation in *Paracoccidioides* using long-read assemblies. Overall, the genus harbors fewer TEs (12–20% of total DNA content) than the related outgroup *Histoplasma* (∼30%; Figure 6A–B). In all isolates, RNA TEs were more abundant than DNA TEs, with RNA:DNA ratios being at least 2.34. Within the *brasiliensis* species group, TE content was relatively stable (12–16% of total DNA) and showed no consistent differences in DNA TE, RNA TE, or their ratios across species pairs. The only notable divergence was observed in *P. lutzii*, which exhibited elevated TE abundance, with TEs comprising ∼19–20% of the genome. This increase was driven primarily by RNA TEs: both *P. lutzii* isolates contained roughly three times more RNA TEs than any other isolate (Table S6, Figure 6A–B, Figure S4).

**FIGURE 6.**
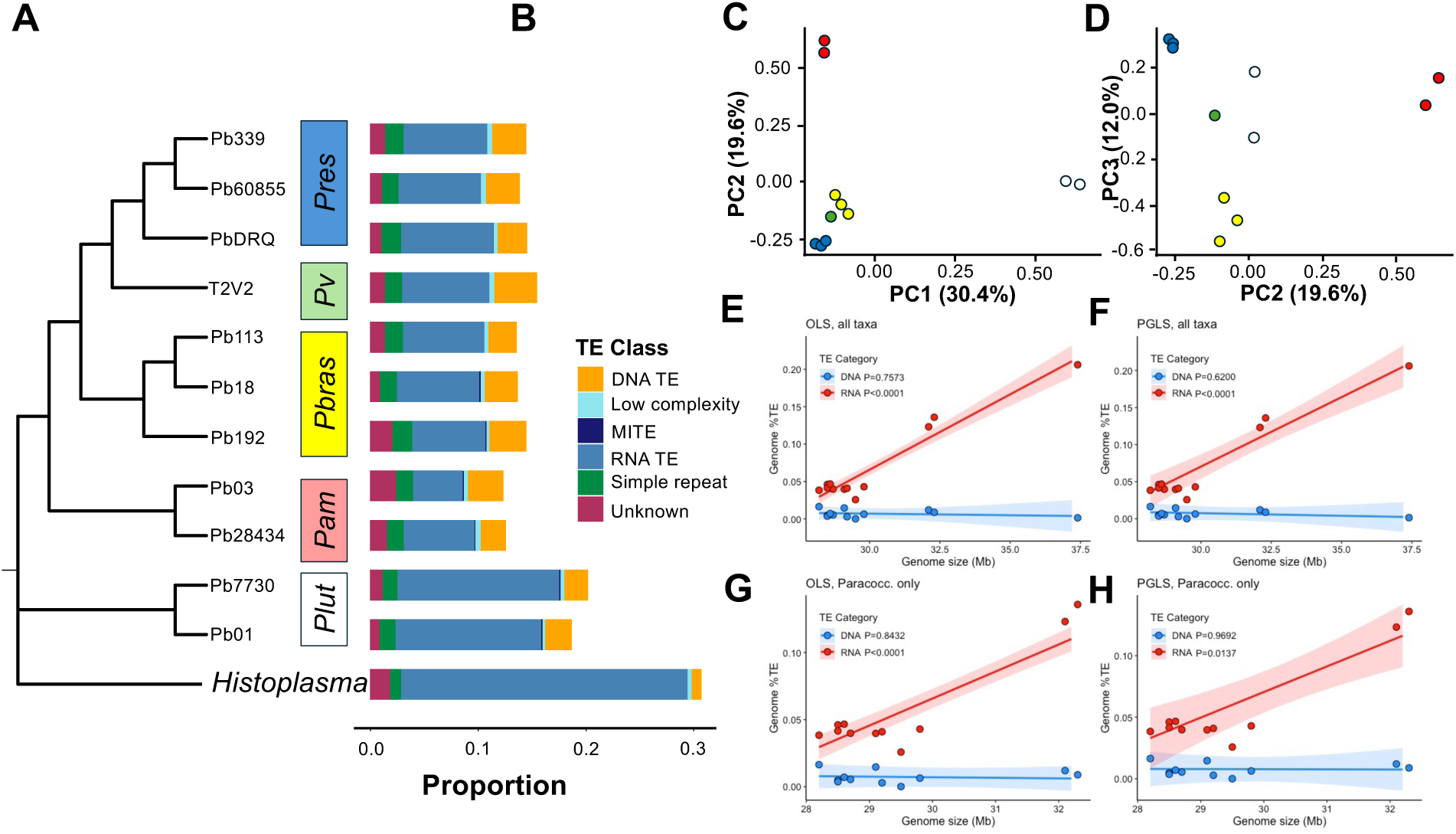
TE evolution. **A** Tree (same as 5A). **B.** Barplots showing the relative frequency of different TEs. The bar plots show DNA TEs, RNA TEs, Low complexity elements, and Miniature Inverted-repeat Transposable Elements (MITEs). Figure S4 shows a similar depicting focusing only on DNA and RNA TEs. **C.** PCA genome. **D.** PCA TEs. Species are labeled by name and color for grouping clarity. Contribution (eigenvalue weight) of each component is displayed in the relevant axis. The Scree plot showing the contribution of other PCS is shown in Figure S5. **E-H.** Correlations of genome size and TE content. Regressions are shown using two methods: ordinary least squares (OLS: **E, F**) and phylogenetic generalized least squares (PGLS: **G, H**). Top panels show regressions including *H. capsulatum*, and the bottom panels show regressions without *H. capsulatum.* Shaded areas indicate 95% confidence intervals.

We also studied the extent of collective differentiation in the TEs of *Paracoccidioides* using PCA. Consistent with previous efforts, we find that the genome wide genetic variation is delimited across species boundaries. The first three components explain 80% of the variance in the dataset (Figure S5). PC1 explains 62.5% of the variance, and differentiates between *P. lutzii* and the species in the *brasiliensis* complex. PC2 primarily distinguishes *P. americana* from the genus. PC3 differentiated *P. brasiliensis, P. venezuelensis,* and *P. restrepiensis* (Figure S6). These results are consistent with previous assessments of genetic diversity in the *Paracoccidioides* genus and indicate that long-read sequencing is able to recapitulate the extent of divergence in the genus. The genetic variance in TEs is also partitioned across species but to a lesser extent that the rest of the genome. When we limit the analysis to TE-only regions, the first three components explain ∼62% of the variance (Figure S5). Similar to the genome-wide results, PC1 differentiates *P. lutzii* from the species of the *brasiliensis* complex, and PC2 and PC3 discriminate among the species of the *brasiliensis* complex (Figure 6C-D).

Next, we studied whether the genome size of *Paracoccidioides* was correlated with TE content. When we included the outgroup, *Histoplasma*, we found significant positive correlations between genome size and RNA TE content using both OLS (P<0.0001, Figure 6E) and PGLS (P<0.0001, Figure 6F). These results held when we excluded *Histoplasma*, which has the largest genome size among our focal taxa (OLS: P<0.0001; PGLS P=0.014; Figure 6G-H). In contrast, we did not find significant correlations between genome size and DNA TE content using either OLS (P=0.757) or PGLS (P=0.620) with *Histoplasma*, or without *Histoplasma* (OLS: P=0.843; PGLS P=0.969; Figure 6E-H).

We examined the two telomeric assemblies (Pb03 and Pb18) to inspect the distribution of TEs along the chromosomes. Both DNA and RNA TEs were more likely to occur at the ends of the chromosomes (Figure 7A). Figure S7 shows the TE distribution for Pb03, and Figures S8-S10 show TE distribution for three highly contiguous, yet not chromosomal, assemblies. Genomic windows that contained DNA TEs were also more likely to contain RNA TEs (Figure 7B). This pattern also holds for the nine highly contiguous genomes and is not dependent on the window size (Figure S11). Notably, the RNA TE content and DNA TE content is not correlated across *Paracoccidioides* isolates, a result that holds before and after phylogenetic corrections (Figure 7C).

**FIGURE 7.**
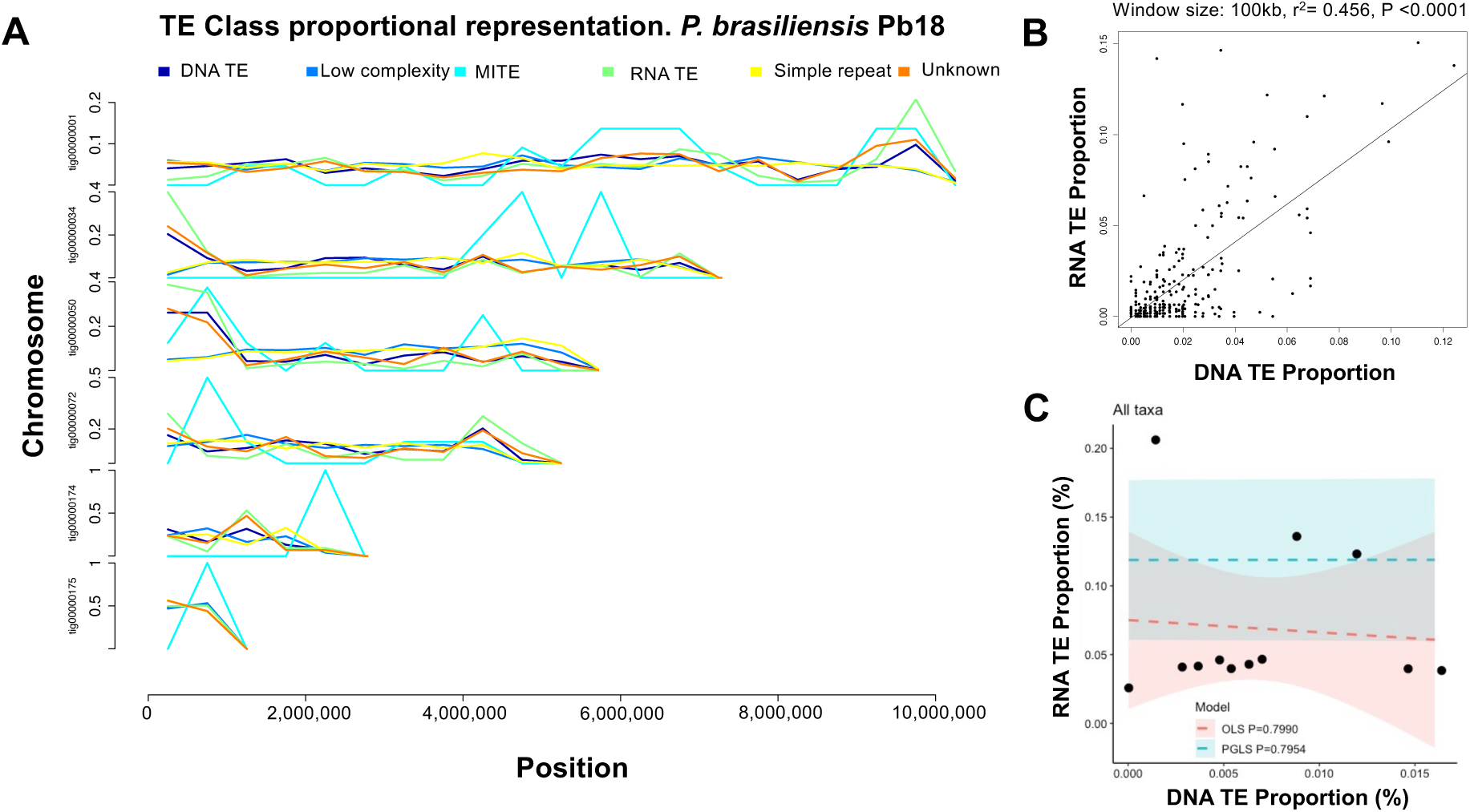
Genomic location of TEs in *Paracoccidioides brasiliensis*. **A.** TE class proportion distribution along the chromosome in a chromosomal assembly of the *P. brasiliensis* genome (Pb18). TE are mostly found at the end of chromosomes. **B.** RNA and DNA tend to concur in the same genome windows (window size =1 × 10^5^). Figure S10 shows a similar analysis with a different window size. **C.** The proportion of DNA TEs and RNA TEs are not correlated across isolates. Blue shows the confidence interval for the ordinary least squares (OLS) regression; red shows the confidence interval for the phylogenetic generalized least squares (PGLS) regression.

### Gene Family Evolution

We studied the expansion and contraction of three gene classes: transcription factors (TF), Carbohydrate-active enzymes (CAZy), and protease/proteinase/proteolytic enzymes (MEROPS database). We describe the results for each of the three classes as follows.

#### Transcription factors

Of the 30 InterPRO TF domains analyzed, most contained only a limited number of copies (1–4), and in nearly all cases the copy number was consistent across isolates (Figure 8A). CAFE5 identified five InterPRO families with statistically significant expansions or contractions: IPR001005 (Myb-like DNA-binding motif TFs), IPR007219 (fungal-specific domain TFs), IPR001138 (fungal Zn(2)-Cys(6) binuclear cluster domain motif TFs), IPR006856 (mating-type protein MAT alpha-1 HMG-box), and IPR001878 (zinc knuckle CCHC motif TFs) (Figure 8A).

**FIGURE 8.**
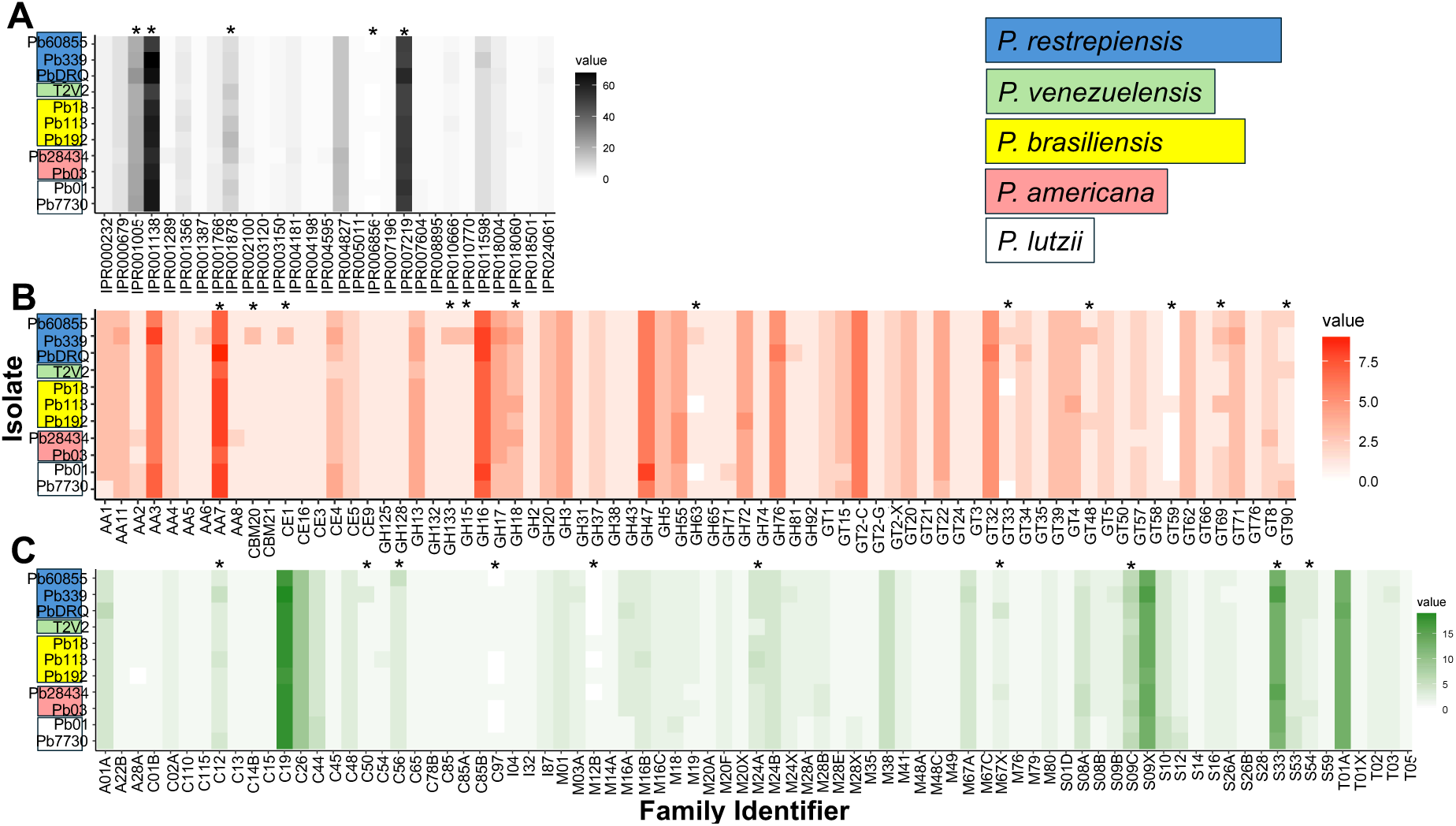
Gene family evolution. The heatmaps on the top panels show the number of hits inferred for each genome. **A.** Transcription factors inferred using the InterPRO database (Paysan-Lafosse et al. 2023; Blum et al. 2025). **B.** Inferred carbohydrate-Active enZYmes database (CAZy, (Cantarel et al. 2009)). **C.** Inferred peptidase/proteinases using the MEROPS database (Rawlings et al. 2010)**. D.** Stacked bars showing the number of CAZy copies per individual. **E.** Stacked bars showing the number of peptidase/proteinases copies per individual.

For IPR001005 (Myb-like DNA binding motif TFs), we found a significant expansion specifically in *P. restrepiensis* strain PrDRQ which contained 26 copies, compared to the 20-23 counts detected in all other *Paracoccidioides* isolates. Similarly, we found a significant family expansion for IPR007219 (Fungal-specific domain TFs) in *P. restrepiensis* strain PbDRQ and *P. lutzii* strain Pb01 (57 and 54 counts respectively), compared to 49-52 counts observed in the remaining *Paracoccidioides* isolates. IPR001138 (Fungal Zn(2)-Cys(6) binuclear cluster domain motif TFs) showed evidence of a gene family contraction in *P. restrepiensis* strain Pb60855 and *P. venezuelensis* strain PvT2V2, reduced to 51 and 50 counts, respectively, compared to the remainder of the *Paracoccidioides* isolates which range from 56-68 counts. IPR006856 (Mating-type protein MAT alpha-1 HMG-Box TFs) showed statistical significance due to the variability in mating type genes present in the pool of *Paracoccidioides* isolates. Mating type genes in *Paracoccidioides* can be present in one of two forms: *Mat1-1* or *Mat1-2* (Torres et al. 2010; Teixeira et al. 2013), so the count ranges for IPR006856 range from 0-1 depending on the class of mating type gene present in the isolate. Lastly, IPR001878 (Zinc Knuckle CCHC motif TFs) have undergone significant gene family evolution which both increases and decreases depending on the isolate. This family showed an expansion in *P. brasiliensis* strains Pb192 and Pb113 (17 and 14 counts respectively), and a contraction in all three *P. restrepiensis* strains: Pb60855, PbDRQ, and Pb339 (8, 9, and 9 counts, respectively). The two *P. americana* isolates, 28434 and Pb03, show a large range of copy numbers with 14 and 9 counts, respectively (Figure 8).

#### Carbohydrate-active enzymes (CAZy)

Twelve carbohydrate-active enzymes underwent gene family size changes (Figure 8B, Figure S12). We identified five glycosyltransferase CAZy enzyme families undergoing significant fold changes across our pool of *Paracoccidioides* isolates (GT90, GT69, GT48, GT33, and GT59). Classes GT90, GT69, and GT48 have each undergone gene family expansions in various isolates, with counts ranging from 1-3 depending on class. GT33 appears to have undergone a family contraction, with two isolates (Pb18 and Pl7730) containing 0 counts of this glycosyltransferase class. The GT59 class shows a unique pattern of being absent in most isolates (0 counts) and appearing in two specific isolates only with a single count (Pb113 and Pb7730).

We detected multiple gene family changes for four glycoside hydrolase classes (GH18, GH63, GH15, and GH133). GH18 ranges from 3-4 counts depending on the isolate, with variance across 3 different *Paracoccidioides* species (*P. restrepiensis, P. brasiliensis, and P. americana*). GH63 has undergone expansion and contraction, with most isolates containing one count of this class. We observed an expansion in isolate Pb339 (2 counts) and the absence of this family (0 counts) in two isolates (Pb113 and Pb01). CAZy enzyme families CE1 (carbohydrate esterase family 1) and CBM20 (carbohydrate-binding module family 20) both showed a unique family expansion specifically only in isolate Pb339, with an increase to three counts of each compared to the single count seen across all other *Paracoccidioides* isolates. In general, Pb339, shows expansions in eight different CAZy enzyme classes (CE1, GT69, GT48, GT33, GH63, GH15, CBM20, and GH133). In particular the types CE1 (carbohydrate esterase), GT48 (glycosyltransferase), GH63 (glycoside hydrolase), and CBM20 (starch-binding), show the largest increases in copy number in this isolate, Figure S12).

#### Proteases, proteinases, and proteolytic enzymes

We identified a total of 10 peptidase-type gene families which underwent gene family evolution (Figure 8C, Figure S13). We found a species-specific expansion event for MEROPS enzyme class S54 (membrane-bound serine endopeptidase) which was expanded in all three isolates of *P. restrepiensis* (Pb-DRQ, Pb339, and Pb01) along with a single isolate of *P. americana* (Pb03). All expanded strains contained three counts of S54-type peptidases, compared to the typical 2 counts observed in all other species. Additionally, *P. restrepiensis* isolates showed family expansions in five MEROPS enzyme classes (C12, C56, S33, C50, and M67X). Similar to CAZy-type enzymes, we found elevated levels of MEROPS class enzymes specifically in isolate Pb339. This *P. restrepiensis* isolate carries elevated counts of multiple MEROPS class enzymes (C12, S33, and C50). This isolate seems to have undergone a very unique large-scale expansion of gene families resulting in Pb339 containing high levels of many enzyme classes (Figure S13).

## DISCUSSION

The evolution of genome structure is a key area of evolutionary genetics that has attracted interest from both evolutionary and clinical biologists, due to the potential impact of genomic rearrangements on fitness (Moyle et al. 2010; Sharma et al. 2021; Araya et al. 2025) and virulence (Jiang and Tyler 2012; Möller and Stukenbrock 2017; Merrikh and Merrikh 2018). In fungal pathogens, for instance, the interplay between effective population size and host interactions can significantly influence the organization of linkage groups within the genome (Dong et al. 2015; Möller and Stukenbrock 2017). Despite growing interest in the genome structure of fungal pathogens, technical limitations have historically hindered progress. While short-read sequencing has been a powerful tool for identifying nucleotide-level variants, it is inherently limited in its ability to detect complex mutations, including chromosomal rearrangements (Lucas Lledó and Cáceres 2013; Kosugi and Terao 2024). Recent advances in long-read sequencing technologies and bioinformatics tools have helped bridge this gap. In this study, we present chromosomal and near-chromosomal genome assemblies for four species of *Paracoccidioides*. Additionally, we provide structural genomic analyses for all known culturable species within the *Paracoccidioides* genus. The implications of these findings are discussed in the sections that follow.

The first implication of our results lies in advancing the understanding of karyotypic evolution in *Paracoccidioides*. Our genome assemblies indicate that the genome size across *Paracoccidioides* species is approximately 30 Mb, with some variation among isolates. These estimates align with previous genome assemblies (Desjardins et al. 2011; Muñoz et al. 2014; Lorenzini Campos et al. 2024) but is considerably smaller than earlier estimates based on pulsed-field gel electrophoresis (Nogueira Cano et al. 1998). Genome sizes within the order Onygenales vary widely—from around 22 Mb in *Ascophaera apis* (Qin et al. 2006) to 75 Mb in *Blastomyces dermatitidis* (Muñoz et al. 2015; McTaggart et al. 2024). A revision of these estimates using chromosomal, or at least highly contiguous assemblies,will reveal whether genome size correlates with ecological lifestyle. Genome size has been proposed to influence diversification rates; however, the evidence supporting this relationship remains inconclusive (Jakob et al. 2004; Kraaijeveld 2010; Kapralov and Filatov 2011; Puttick et al. 2015).

The two telomeric assemblies reveal a dynamic chromosomal structure within the *Paracoccidioides* genus. The species *P. americana* and *P. brasiliensis* diverged between 0.117 and 0.714 million years ago, depending on the molecular model used to estimate divergence time (Teixeira et al. 2009b; Turissini et al. 2017; Mavengere et al. 2020). Although both isolates possess five chromosomes, the chromosomes are not entirely homologous, despite the presence of large syntenic blocks. This discrepancy is the result of one chromosomal fusion and one fission event. Future studies with additional chromosome-level assemblies will be needed to determine whether these rearrangements are fixed species-specific traits or represent polymorphic variation within populations. The highly contiguous genomes we present suggest the latter, but answering this question completely will require approaches that go beyond simply identifying chromosomal changes and instead assess their allelic frequencies within and across populations.

We also detected more granular rearrangements. We focused on the two species that had chromosomal assemblies but that does not mean the other species are not of interest. We detected one highly supported tandem duplication in *P. americana*. Of note, this 90kb duplication contains seven genes. Three of these genes make part of the siderophore biosynthetic pathway: *sid3, sid4,* and *sid1*. *sid3* is orthologous to *sidF* in *Aspergillus fumigatus*, an N^5^-transacylase involved in the synthesis of fusarinine and triacetylfusarinine. *ΔsidF* mutants in *A. fumigatus* show a reduced conidiation rate on iron-limiting conditions, partial sensitivity to oxidative stress, and significantly reduced virulence on neutropenic mice (Schrettl et al. 2007). *sid4* shows homology to *sidI* in*Aspergillus terreus*, a mevalonyl-CoA ligase. In *A. fumigatus*, *ΔsidI* mutants are susceptible to iron starvation and oxidative stress and show reduced virulence in neutropenic mice (Yasmin et al. 2012). In *Histoplasma*, *sid3* and *sid4* belong to a cluster of genes (93 genes) that are upregulated in response to dbcAMP but to date no effort has genetically evaluated their contribution to virulence.

Of these three genes, the best characterized is *sid1*. *sid1*, encodes a L-ornithine monooxygenase, the enzyme responsible for the first step in siderophore biosynthesis in multiple fungal pathogens ((Mei et al. 1993; Eisendle et al. 2003) reviewed in (Howard 2004; Nyilasi et al. 2005)). The ortholog of *sid1* in *Aspergillus fumigatus, sidA,* is essential for virulence (Schrettl et al. 2004). In *Histoplasma*, *sid1* is part of a cluster of co-upregulated genes under low-iron conditions, all involved in siderophore synthesis, secretion, and utilization (Hwang et al. 2008). The disruption of *sid1* by allelic replacement impaired growth in low-iron conditions and eliminated siderophore production in *H. ohiense*. *sid1*-deficient strains showed reduced growth in murine bone marrow-derived macrophages and were attenuated in a mouse infection model (Hwang et al. 2008; Brechting et al. 2023 July 11). In *P. brasiliensis*, RNA-silencing mutants of *sidA* (equivalent to *sid1*) show decreased siderophore biosynthesis in iron poor conditions and reduced virulence to beetle (*Tenebrio molitor*) model (Silva et al. 2020). Gene duplication constitutes an important evolutionary phenomenon because duplicated copies can acquire new functions, new expression domains, or specialize in a particular function (reviewed in (Innan and Kondrashov 2010; De Smet and Van de Peer 2012; Conant et al. 2014)). While previous studies have measured the virulence of *P. americana* Pb03 and have found that this isolate actually shows lower virulence than other *Paracoccidioides* isolates (Carvalho et al. 2005), an assessment of the effect of duplicating these genes on virulence remains to be examined with a larger sample that takes into account the rest of the genetic background.

Our studies also revealed evolutionary patterns of the mtDNA genome in *Paracocccioides*. First, we find that the mtDNA shows complete synteny within the *Paracoccidioides* genome. The inferred gene order of the mtDNA genome is consistent with previous reports (Cardoso et al. 2007; Misas et al. 2020). Second, the expansion of phylogenetic studies to the entirety of the mitochondrial genome also solves a conundrum in evolutionary studies of *Paracoccidioides*. Multilocus sequenced typing had reported the existence of two mitochondrial lineages within *Paracoccidioides* which was reported as inconsistent with nuclear assessments of genetic structure within the genus (Salgado-Salazar et al. 2010; Turissini et al. 2017). Mitochondrial capture is a common phylogenetic phenomenon in plants and animals but so far remains mostly unstudied in fungi. The case in *Paracoccidioides*, both the topology and the length of the tree indicate that mtDNA has been introgressed within species and is likely that *P. venezuelensis* and *P. lutzii* have hybridized in Venezuela. A more comprehensive assessment with more isolates, and full genome assemblies, will determine whether there has been a full mtDNA genome replacement or whether there are different mtDNA haplotypes segregating. These results are significant because surveys for introgression in the nuclear genome of *Paracoccidioides* suggest little evidence of haplotypes derived from admixture between species (Mavengere et al. 2020). On the other hand, the mitochondrial genome does indicate gene exchange in the recent past. One possibility that could explain these results is incomplete lineage sorting of the mtDNA genomes. If different forms of the mtDNA existed before speciation, then fixation of the different haplotypes would not necessarily reflect the evolutionary history of the clade, thus leading to a different topology (Good et al. 2008; Marková et al. 2013; Good et al. 2015; Andersen et al. 2021). This hypothesis is unlikely given the short tip-to-tip distance of the mtDNA tree compared to that of the nucDNA. This pattern is the opposite of the expectation of tree length under an mtDNA ILS scenario in which the mtDNA would have a deeper divergence than the nucDNA tree. A second potential explanation for this pattern is that the nuclear genome carries hybrid incompatibilities which in turn can lead to the rapid purging of introgression after hybridization. If incompatibilities are evenly distributed along the genome, this would lead to a rapid extinction of the haplotypes of one of the hybridizing species (Matute et al. 2020). This decline is expected to be more precipitous when the hybridizing species are highly diverged (Dagilis and Matute 2023). In this scenario, the nuclear genome would revert rapidly to that of one of the parents but if there are no mito-nuclear incompatibilities, the mtDNA can be exchanged readily among species. This hypothesis is out of the scope of what we can test because it would require experimental crosses between isolates of different species, and assessments of hybrid fitness, a challenging task as the sexual stage of *Paracoccidioides* has never been produced in laboratory conditions.

We also measure the extent of variation in TE content in the genus *Paracoccidioides*. TEs have been proposed to be an important player in generating genome variability in fungi (reviewed in (Sauters and Rokas 2025)). Our studies revealed that RNA TE, but not DNA TE, content is one of the underlying causes of the difference in genome size among isolates of *Paracoccidioides*. Similar patterns have been observed in many taxa (Lee and Kim 2014; Cong et al. 2022; Dai et al. 2022) but the positive correlation is not universal (Zuo et al. 2023). Our TE survey did reveal at least two aspects of TE evolution in *Paracoccidioides*. First, the TE content is lower than that in the *Histoplasma* genome (Voorhies et al. 2022) and *Blastomyces* (McTaggart et al. 2024). Similar to *Histoplasma* (Voorhies et al. 2022) and *Ganoderma* (Wang et al. 2023), *Paracoccidioides* TEs are located towards the end of chromosomes, a pattern that does not seem to be universal for fungi. Second, *P. lutzii* contains more RNA TEs than the rest of *Paracoccidioides*, indicating parallel accumulation of TEs in *Histoplasma* and *P. lutzii* or a reduction in TE content in the species of the *brasiliensis* species complex. A more comprehensive phylogenetic survey will reveal the patterns of evolution of TEs in individual species. Our sample size does not allow us to determine what is the tempo and evolutionary mode of TE evolution in the genus but comparative phylogenetic methods have the potential to discern how these accumulations have taken place.

Our results also reveal changes in gene family size within the *Paracoccidioides* genus. Carbohydrate-active enzymes (CAZy), for instance, vary among *Paracoccidioides* isolates. Glycosyltransferases, a CAZy family often associated with virulence in plant pathogens (e.g., (King et al. 2017; Deng et al. 2021; Blandenet et al. 2022), show notable differences. In the yeast *Metschnikowia bicuspidata*, a crustacean pathogen, four of the five CAZy classes (GHs, GTs, CEs, and CBMs—but not AAs) have undergone rapid and extensive expansion (Shi et al. 2023). In human pathogens, N-linked mannosylation is essential for cell adhesion and virulence in *Candida* (reviewed in (Martínez-Duncker et al. 2014)),often contributing to biofilm formation and serving as a major virulence factor in many pathogenic fungi (Danchik and Casadevall 2021). The observed expansion of peptidases in *Paracoccidioides* may provide insight into its life cycle and pathogenic potential. Similar to other animal fungal pathogens (e.g., (Li et al. 2012; Gutierrez-Gongora and Geddes-McAlister 2022)*, Paracoccidioides* may have experienced expansions in gene families associated with mammalian parasitism.

The generation of genome alignments from long-read assemblies will facilitate the identification of structural variants and the investigation of non-point mutations underlying phenotypic differences among pathogens, including their potential roles in virulence. Our results indicate that such variants are present in *Paracoccidioides* and are polymorphic within species. Systematic assessments of their allele frequencies and trajectories over time could reveal whether these changes have been shaped by natural selection.

## DATA AVAILABILITY

All scripts used for the analyses, together with the reference annotation files and the consensus TE file used in this study, are available at https://github.com/Dogrinev/

## ACKNOWLEDGEMENTS

Thanks to the Núcleo de Coleção de Micro-organismos, Instituto Adolpho Lutz, São Paulo, Brazil and Coleção de Fungos Patogênicos, INI/Fiocruz, Rio de Janeiro, Brazil. We are grateful to the Matute lab for comments in previous versions of this manuscript.

